# C5aR1 antagonism alters microglial polarization and mitigates disease progression in a mouse model of Alzheimer’s disease

**DOI:** 10.1101/2022.05.18.492363

**Authors:** Angela Gomez-Arboledas, Klebea Carvalho, Gabriela Balderrama-Gutierrez, Shu-Hui Chu, Heidi Yahan Liang, Nicole D. Schartz, Purnika Selvan, Tiffany J. Petrisko, Miranda A. Pan, Ali Mortazavi, Andrea J. Tenner

## Abstract

Multiple studies have recognized the involvement of the complement cascade during Alzheimer’s disease pathogenesis; however, the specific role of C5a-C5aR1 signaling in the progression of this neurodegenerative disease is still not clear. Furthermore, its potential as a therapeutic target to treat AD still remains to be elucidated. Canonically, generation of the anaphylatoxin C5a as the result of complement activation and interaction with its receptor C5aR1 triggers a potent inflammatory response. Previously, genetic ablation of C5aR1 in a mouse model of Alzheimer’s disease exerted a protective effect by preventing cognitive deficits. Here, using PMX205, a potent, specific C5aR1 antagonist, in the Tg2576 mouse model of Alzheimer’s disease we show a striking reduction in dystrophic neurites in parallel with the reduced amyloid load, rescue of the excessive pre-synaptic loss associated with AD cognitive impairment and the polarization of microglial gene expression towards a DAM-like phenotype that are consistent with the neuroprotective effects seen. These data support the beneficial effect of a pharmacological inhibition of C5aR1 as a promising therapeutic approach to treat Alzheimer’s disease. Supportive of the safety of this treatment is the recent FDA-approval of another other C5a receptor 1 antagonist, Avacopan, as a treatment for autoimmune inflammatory diseases.

**One Sentence Summary:** C5aR1 antagonist shifts microglial gene expression toward neuroprotection.

## INTRODUCTION

Alzheimer’s disease (AD) is the most prevalent neurodegenerative disease of the elderly. Currently, over 6 million people suffer from AD in the US alone, and the affected number of people is expected to rise to 13 million by 2050(*1*). AD is neuropathologically characterized by the extracellular accumulation of Aβ in amyloid plaques and hyperphosphorylated tau in neurofibrillary tangles. The appearance of dystrophic neurites, synaptic loss, neuronal death and finally, cognitive deficits are also hallmarks of Alzheimer’s disease(*2*). Evidence from AD patients, including GWAS studies, strongly point to an inflammatory response as a key mediator in the onset and progression of the disease(*3–5*). In fact, GWAS studies have identified more than 70 genetic loci associated with a high risk to develop Alzheimer’s disease and interestingly, among those genes, two complement genes have been confirmed as significantly associated with AD(*6*).

The complement system (C’) is a very important component of the innate immune system that plays a key role in host defense to ultimately maintain brain homeostasis. Several studies have identified multiple complement proteins, such as C1q, C3 or C4, co-localizing with amyloid plaques not only in mouse models of AD but also in AD patients(*7–9*), suggesting a role of the complement system in AD. In fact, during Alzheimer’s pathogenesis, the complement system can be activated by both, fibrillar Aβ and hyperphosphorylated tau, ultimately leading to the production of the activation fragment C3a and C5a that interact with its cellular receptor C3aR and C5aR1, triggering a potent inflammatory response(*10, 11*). Whether complement activation in the Alzheimer’s brain is beneficial or detrimental is still controversial, as we have previously reported a neuroprotective role of C1q(*12, 13*) but we and others have also demonstrate a beneficial effect of a genetic ablation of C1q, C3 and CR3(*14–16*). Altogether, these data suggest that a complete blockage of the complement cascade might not be the best therapeutic approach for AD; instead, a strategic and very specific therapeutic targeting might be required for this neurodegenerative disorder, such as a pharmacological inhibition of C5a-C5aR1 signaling that would preserve the beneficial effects of the upstream components of the complement cascade while preventing downstream detrimental effects.

Previous studies in our lab using the Arctic48 mouse model of AD(*17*) showed that genetic ablation of C5aR1 completely prevented the loss of neurite complexity and cognitive deficits associated with Alzheimer’s disease; whereas overexpression of C5a by a transgene under the GFAP promoter in the same mouse model seems to accelerate hippocampal-dependent spatial memory deficits(*18, 19*). In addition, pharmacological inhibition of C5a-C5aR1 signaling with a C5aR1 antagonist (PMX205) in two different mouse models of Alzheimer’s disease (the Tg2576 and the 3xTgAD mouse models) resulted in a significant reduction of the neuropathology associated with AD as well as an improvement of memory and cognition(*20*), further supporting the targeted inhibition of C5aR1 as a potential therapeutic target for AD. PMX205 is a cyclic hexapeptide that acts as a potent insurmountable inhibitor of the C5a receptor 1 (C5aR1)(*21*). Pharmacokinetics studies on PMX205 have shown its high oral bioavailability and its efficiency in entering the intact CNS(*21*). Furthermore, an oral daily dosing of PMX205 for long periods of time (where PMX205 was administered through the drinking water) showed maintained levels of PMX205 in the CNS and proved its efficiency to block C5a-C5aR1 signaling in the brain(*20–22*). We and others have previously reported several beneficial effects of this C5aR1 antagonist, not only in Alzheimer’s disease but also in amyotrophic lateral sclerosis and spinal cord injury(*20, 23, 24*). More importantly, PMX53 (which is considered the parent molecule of PMX205) and Avacopan (a small-molecule FDA-approved C5a receptor1 antagonist) have already been proven to be safe in human clinical trials for autoimmune diseases(*25–27*), suggesting that if beneficial effects of PMX205 are demonstrated, it could have an accelerated path to human clinical trials.

Despite the discovery of PMX205 more than 20 years ago and its reported beneficial effects in Alzheimer’s disease models(*20*), the specific mechanism that might be driving the improvement in cognition is still not know. Here, we used the Tg2576 mouse model of Alzheimer’s disease and treated them with PMX205 at the onset of the amyloid pathology to further determine the effects of this C5aR1 antagonist on microglial cells. The results presented in this study demonstrated a neuroprotective effect of PMX205, which rescues the excessive intracellular synaptic proteins and pre-synaptic loss associated with Alzheimer’s disease. This finding seems to be linked to the reduction of a unique microglial subpopulation associated with synaptic pruning in the PMX205 treated mice. Interestingly, we also show here that blocking C5a-C5aR1 signaling in the Tg2576 mouse model of AD results in the increase of the DAM2 gene-expressing microglial subpopulation, suggesting that PMX205 might be blocking disease enhancing events and facilitating a disease mitigating phenotype on microglial cells. Our data further supports the use of C5aR1 antagonists as potential therapeutic targets to treat or slow the progression of Alzheimer’s disease.

## RESULTS

### PMX205 treatment reduces plaque pathology in the Tg2576 mouse model of Alzheimer’s disease

C5aR1 antagonist, PMX205, was administered to WT and Tg2576 mice in their drinking water for 12 weeks. The starting point of the treatment was set up at 12 months of age, coincident with the onset of the amyloid pathology in this Alzheimer’s mouse model (**Sup Fig. 1A**). During the whole duration of the treatment, mice and water bottles were weighted weekly in order to determine possible weight loss negatively associated with the treatment as well as the dose of PMX205 taken by each individual mouse. Overall, WT mice weight was higher than AD mice, as expected. No weight loss was observed in any of the experimental groups during the course of the treatment (**Sup Fig. 1B**), indicative of a lack of a toxic effect of PMX205. All mice consumed similar amounts (ml) of vehicle (water) or PMX205 (diluted in water) per week (**Sup Fig. 1C**) and no significant differences were observed across the experimental groups. The dose range of PMX205 each mice received was around ~ 2-8 mg/kg/day for both, WT and Tg2576 mice. These results are consistent with previous studies in our lab(*20*).

We previously reported a decrease in ThioS+ fibrillar amyloid plaques in the Tg2576 mice treated with PMX205(*20*). Here we stained coronal sections from all four experimental groups with ThioS and OC, for the characterization of the Aß plaques (**Sup Fig. 2A**). As expected, no plaques were detected in WT mice (data not shown), so we excluded them from the quantification. After 12 weeks of treatment with PMX205, Tg2576 mice showed a significant decrease in fibrillar amyloid load (measured by ThioS – Field Area %) when compared to vehicle treated mice (~ 50% and ~ 58% reduction in the hippocampus and cortex, respectively). In addition, we observed a strong reduction in the number of ThioS+ plaques but not in their size, in both regions analyzed (**Sup Fig. 2 B1-B3 and C1-C3**). To continue the characterization of the amyloid pathology, we used the OC antibody, that stains not only for amyloid fibrils but also for fibrillar oligomers (but not prefibrillar oligomers). Our results showed reduced levels of OC % Field area (reduction of ~ 65% and ~ 53% in the hippocampus and cortex, respectively) in the PMX205 treated Tg2576 mouse model (**Sup Fig. 2 B4 and C4**). Similar to ThioS results, the number of OC+ plaques was substantially affected by C5aR1 inhibition (~ 64% and ~ 48% reduction in hippocampus and cortex, respectively), but no changes were found in Aß plaque size (**Sup Fig. 2 B5-B6 and C5-C6**), suggesting that PMX205 might have an effect in the initial plaque deposition stages, but once the amyloid plaques are formed, inhibition of C5a-C5aR1 signaling does not have an effect on them. Positive correlation of ThioS (FA%) and OC (FA%) were found in the hippocampus and cortex of Tg2576 mice (**Sup Fig. 2 B7 and C7**).

### Pharmacological inhibition of C5aR1 significantly affects the expression of several microglial activation markers

C5a receptor 1 is highly expressed upon injury or disease in humans and mouse models including AD(*28–31*). Here, to investigate changes in microglial cells in the brain of WT and Tg2576 mice treated with vehicle or PMX205, coronal sections of all experimental groups were stained using different microglial markers: Iba1, CD68, Cd11b and Cd11c (**Fig. 1 A-B**). Quantitative analysis of the hippocampus and cortex of all four experimental groups revealed significant increases in Iba1, CD68, CD11b and CD11c (FA%) between Tg2576 and WT vehicle treated mice. This effect was counteracted by the presence of a C5aR1 antagonist in the Tg2576 mice (when compared to Tg2576-vehicle mice) for all markers analyzed (**Fig. 1 C1-C4** and **D1-D4**). Overall, our results showed a hippocampal reduction of ~ 38% (Iba1), ~ 23% (CD68), ~ 65% (Cd11b) and ~ 82% (Cd11c) when comparing Tg2576-PMX205 to Tg2576-vehicle treated mice. The reduction in microglial field area percent was, in all cases, positively correlated with amyloid levels in both regions analyzed (**Fig. 1 C5-C8** and **D5-D8**), suggesting that the reduction in microglial coverage in the hippocampus and cortex is a direct response to the reduction in amyloid load in PMX205 treated AD mice. Our results also showed a significant increase in the number of Iba1+ cells (measured as the number of Iba1+ cells per mm^2^) in the hippocampus of Tg2576 vehicle treated mice when compared to WT mice (**Fig. 1 C9**), indicative of an AD-related gliosis.

**Fig 1.**
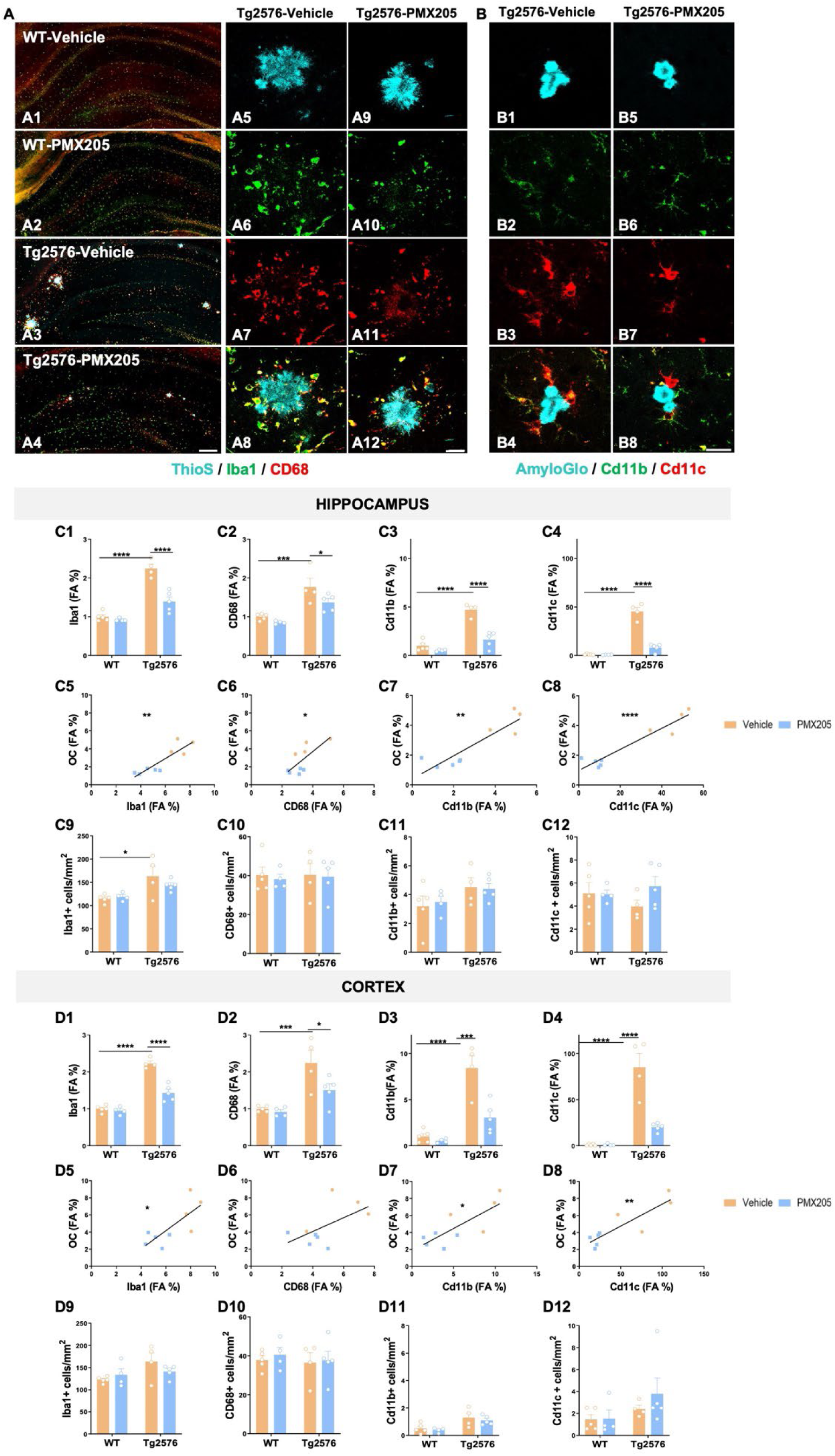
Treatment with PMX205 reduced the expression of several microglial activation markers. **(A-B)** Representative images of the hippocampus of WT-vehicle **(A1),** WT-PMX205 **(A2),** Tg2576-vehicle **(A3)** and Tg2576-PMX205 **(A4)** stained with ThioS (cyan), Iba1 (green) and CD68 (red). Inserts **(A5-A12)** shows representative higher magnification pictures of microglial cells surrounding amyloid plaques in the hippocampus of Tg2576 vehicle/PMX205. **(B1-B8)** Representative images of Cd11b (green) and Cd11c (red) positive microglial cells surrounding Aß deposits in the hippocampus of Tg2576-vehicle **(B1-B4)** or Tg2576-PMX205 **(B5-B8). (C-D)** Field Area (%) quantification of microglial markers in the hippocampus **(C1-C4)** and cortex **(D1-D4)** of Tg2576 treated with PMX205 when compared to the control group. Microglia showed a positive correlation with amyloid plaques in both, the hippocampus **(C5-C8)** and cortex **(D5-D8)** of Tg2576 mice. Quantitative analysis of microglial density **(C9-C12; D9-D12)** measured as number of Iba1, CD68, Cd11b or Cd11c+ cells per mm^2^ in the hippocampus and cortex. Data are shown as Mean ± SEM and normalized to control group (WT-vehicle). Statistical analysis used a 2-way ANOVA and Pearson’s correlation test. Significance is indicated as * p < 0.05; ** p < 0.01; *** p < 0.001 and **** p < 0.0001. Pearson’s R^2^ = 0.78 (C5); 0.58 (D5); 0.54 (C6); 0.32 (D6); 0.79 (C7); 0.61 (D7); 0.90 (C8) and 0.76 (D8). 3 sections/mouse and n = 4-5 mice per group. Scale bar: A1-A4: 200 µm; A5-A6 and B1-B2: 20 µm.

In the cortex of these mice, the number of Iba1+ cells followed a similar pattern of increase in the Tg2576 (vs WT) but did not reach the statistical significance (**Fig. 1 D9**). Additionally, in both regions analyzed, PMX205 showed a trend towards a decrease of the total number of Iba1+ cells in AD mice. Our results regarding the number of CD68 and Cd11b positive cells showed no significant differences among any of the experimental groups (**Fig. 1 C10-C11** and **D10-D11**). Although no significant changes were found regarding the number of Cd11c+ cells, it was interesting that Tg2576-PMX205 showed a trend towards increased number of Cd11c+ cells, when compared to Tg2576-vehicle mice (**Fig. 1 C12**) in the hippocampus. These results matched our findings with single cell RNAseq detailed later in this manuscript.

To investigate whether blockage of C5a-C5aR1 signaling would have a direct effect on amyloid-ß phagocytosis by microglial cells and microglia recruitment around plaques, the amount of amyloid-ß that was detected within the microglial lysosomes in the Tg2576-PMX205 mice was compared to Tg2576-vehicle mice (**Fig. 2A**). No significant changes between the groups were detected. In addition, no differences were found when we assessed microglial engagement around Aß plaques when comparing PMX205 treated AD mice to vehicle treated AD mice (**Fig. 2B**). Taken together, these results suggest that the reduction in the total amount of amyloid plaques by PMX205 is not a direct effect of increased Aß phagocytosis by microglial cells. Furthermore, as we previously described before, the reduction in the number of plaques but not in their size, further support an effect of C5aR1 on the initial plaque deposition stages.

**Fig 2.**
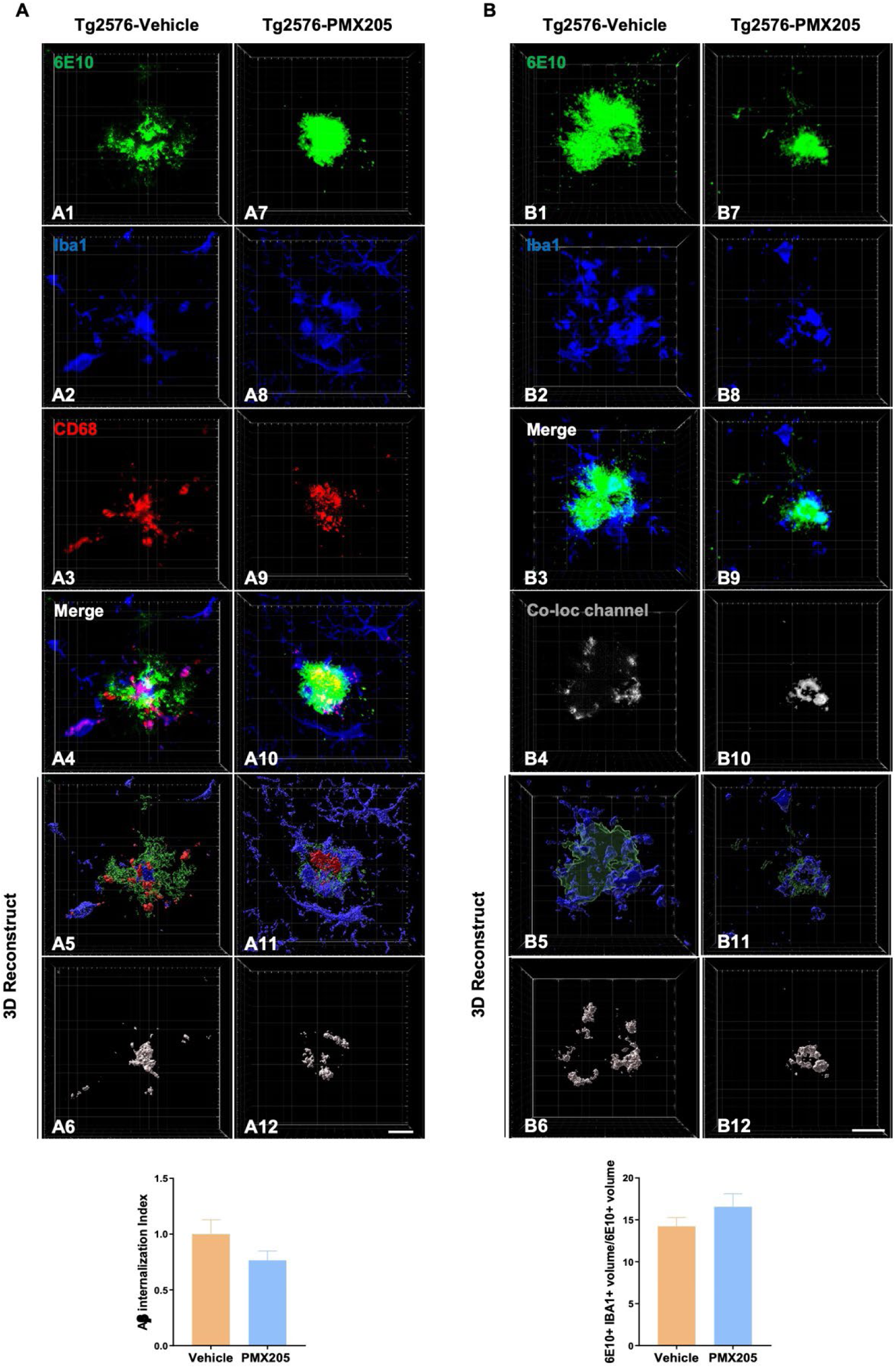
Impact of PMX205 on microglia-plaque interaction in the Tg2576 mouse model of Alzheimer’s disease. **(A)** Representative immunohistochemical images of amyloid plaques 6E10: green), microglial cells (Iba1: blue) and microglial lysosomes (CD68: red) in the Tg2576-vehicle **(A1-A6)** and Tg2576-PMX205 **(A7-A12).** Imaris 3D-reconstruction images showed a trend towards reduced internalization of Amyloid-ß by microglial cells **(A6 and A12)** in the Tg2576-PMX205 mice when compared to the vehicle treated group. **(B)** Representative images of amyloid plaques (6E10: green) and microglial cells (Iba1: blue) in the Tg2576-vehicle **(B1-B4)** and Tg2576-PMX205 **(B7-B10).** Tridimensional reconstruction images **(B5-B6 and B11-B12)** and quantification showed no difference in the engagement of microglial cells around amyloid plaques between the PMX205 treated group and the controls. Data are shown as Mean ± SEM and normalized to control group (Tg2576-PMX205). Statistical analysis used a two-tailed t-test. 15 amyloid plaques/mouse and n = 4-5 per group. Scale bar: 10 µm.

### Treatment with a C5aR1 antagonist exerts neuroprotective effects in the Tg2576 mouse model of Alzheimer’s disease

Given our previously reported results, where pharmacological inhibition of C5a-C5aR1 signaling or genetic ablation of C5aR1 significantly rescue memory deficits in two mouse models of Alzheimer’s disease(*18, 20*), we further explored the potential beneficial effect of PMX205 on the appearance of dystrophic neurites (swollen dendrites and/or swollen axons that appears surrounding neuritic amyloid plaques) and microglial synaptic pruning. Coronal sections from all four experimental groups were stained with ThioS, OC and Lamp1, to detect both amyloid plaques and dystrophic neurites. As expected, neither ThioS+/OC+ plaques nor Lamp1+ dystrophic neurites accumulate in WT mice (images not shown), so WT groups were excluded from the quantification analysis. In contrast, AD mice (Tg2576) showed a strong accumulation of dystrophic neurites surrounding Aß plaques within the hippocampus and cortex (**Fig. 3A**). Interestingly, Lamp1 quantification on the Tg2576-vehicle versus Tg2576-PMX205 reveals a significant reduction of the total amount of dystrophic neurites due to the inhibition of C5aR1 in both regions analyzed (~ 48% and ~ 57% reduction in the hippocampus and cortex respectively) (**Fig. 3 A3** and **A6**). However, the ratio of dystrophic neurites (FA%) per ThioS+ plaques (FA%) did not change among the two groups, suggesting that amyloid plaques are eliciting the same toxicity/damage independently of C5aR1 signaling (**Fig. 3 A4** and **A7**). In line with this, we also observed a significant positive correlation of Lamp1 and amyloid plaques in both regions analyzed (**Fig. 3 A5** and **A8**).

**Fig 3.**
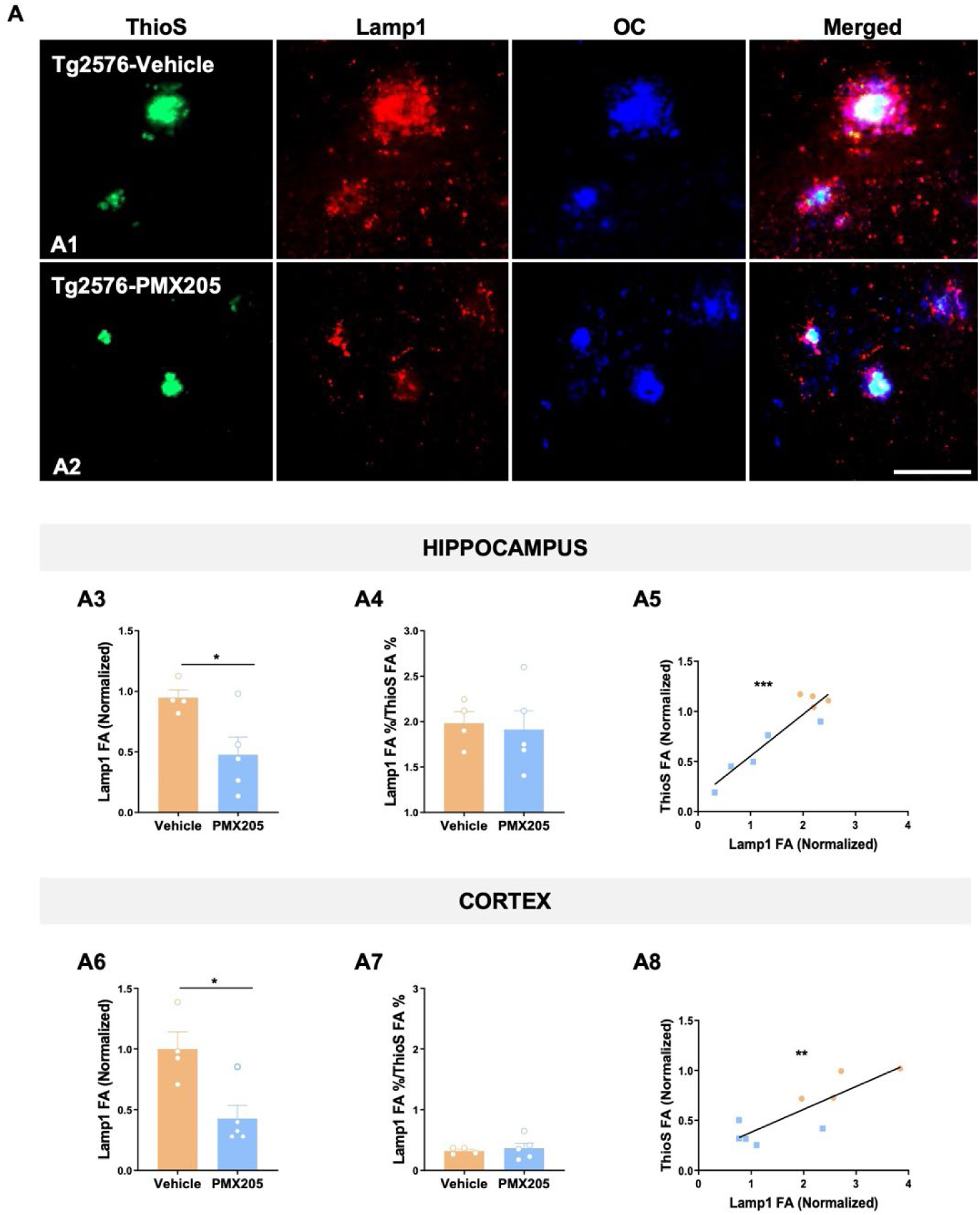
Treatment with PMX205 showed a beneficial effect in reducing the dystrophic neurites pathology on the Tg2576 Alzheimer’s mice. **(A)** Representative images of amyloid plaques (ThioS: green), dystrophic neurites (Lamp1: red) and protofibrillar amyloid species (OC: blue) in the hippocampus of Tg2576 mice treated with vehicle **(A1)** or PMX205 **(A2).** Quantitative analysis of Lamp1 showed a significant reduction in the appearance of dystrophic neurites in mice treated with PMX205 in both hippocampus and cortex (**A3 and A6).** Analysis of the ratio Lamp1/ThioS showed no difference between vehicle and PMX205 treated mice **(A4 and A7).** Pearson’s correlation test showed a positive correlation of Lamp1 with ThioS+ plaques in the hippocampus (R2 = 0.87; **A5**) and cortex (R2 = 0.71; **A8**). Data are shown as Mean ± SEM and normalized to control group (Tg2576-vehicle). Statistical analysis used a two-tailed t-test and Pearson’s correlation test. Significance is indicated as * p < 0.05; ** p < 0.01; and *** p < 0.001. n = 4-5 per group. Scale bar: A1-A2: 100 µm.

Synaptic loss is an important hallmark of Alzheimer’s disease and best correlates with the cognitive decline detected in Alzheimer’s patients(*32*). Furthermore, it has been suggested by our lab and others that excessive synapse loss is due to an excessive complement-mediated synaptic pruning(*9, 33–36*). To explore microglial synaptic pruning, we evaluated Iba1 (microglial cells), CD68 (lysosomes) and VGlut1 (pre-synapse) in all four experimental groups by IHC. High resolution images of 15 individual microglial cells per mouse were acquired (**Fig. 4**) and quantification of the total amount of pre-synaptic puncta (VGlut1+) and engulfed pre-synapses by microglial cells was carried out. Our results demonstrated a significant loss of VGlut1+ puncta in CA3 region in the Tg2576-vehicle when compared to WT littermates (**Fig. 4 A6**). Furthermore, when assessing the synaptic engulfment by microglial cells we observed increased engulfment of VGlut1 in the Tg2576-vehicle (vs WT-vehicle) and a significant decrease of microglial synaptic engulfment in the Tg2576-PMX205 when compared to vehicle treated AD mice (**Fig. 4 A5**). Interestingly, the Tg2576-PMX205 mice showed a rescue of the excessive loss of VGlut1 puncta when compared to Tg2576 vehicle treated (**Fig. 4 A6**) in the CA3 region of the hippocampus. These results point to an excessive microglial synaptic pruning, in our Alzheimer’s mouse model that could be the main cause of the synaptic loss observed in these mice. These observations correlate with previous results in our lab where we observed that the cognitive deficits in the Tg2576 mouse model of Alzheimer’s disease were rescued by PMX205 administration(*20*). Together, our results demonstrated a neuroprotective and beneficial effect of blocking C5a-C5aR1 signaling in the Tg2576 mouse model of Alzheimer’s disease.

**Fig 4.**
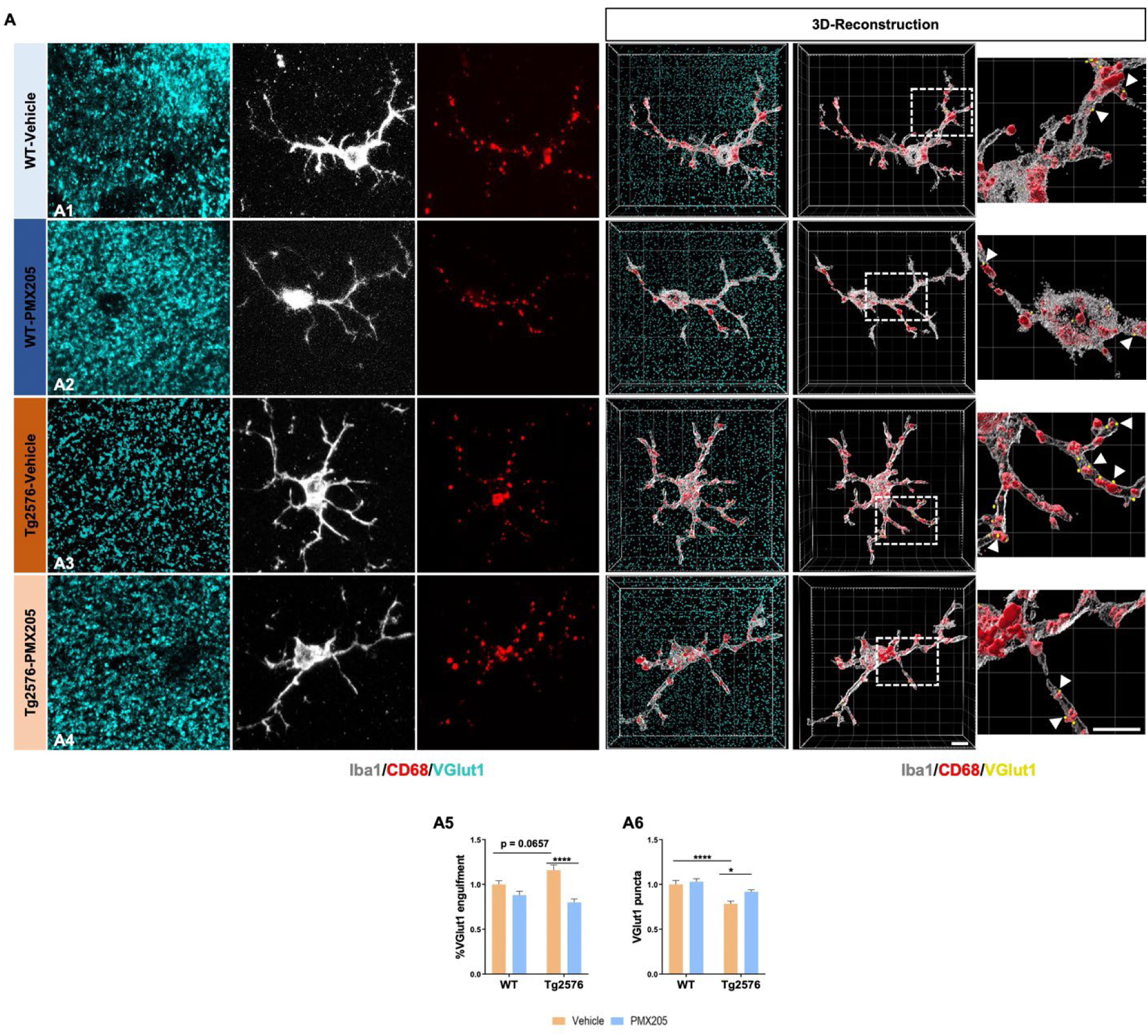
Pharmacological inhibition of C5aR1 rescues VGlut1 loss and reduces synaptic engulfment by microglial cells in the Tg2576 mouse model of AD. **(A)** Representative confocal images and tridimensional reconstruction of individual microglial cells (Iba1: gray), pre-synapses (VGlut1: cyan) or internalized pre-synapses (VGlut1: yellow and arrowheads) and microglial lysosomes (CD68: red) in the CA3 hippocampal region of WT and Tg2576 treated with vehicle or PMX205 **(A1-A4).** Quantitative analysis of the number of pre-synapses internalized by microglial cells revealed a decrease in the synaptic phagocytosis by microglial cells in the Tg2576-PMX205 mice when compared to Tg2576-vehicle **(A5).** VGlut1 puncta quantification showed a strong reduction of pre-synapses in the Tg2576 mice when compared to WT mice; that was restored in the Tg2576 mice treated with PMX205 vs vehicle treated mice **(A6).** Data are shown as Mean ± SEM and normalized to control group (WT-vehicle). Statistical analysis used a 2-way ANOVA. Significance is indicated as * p < 0.05; and **** p < 0.0001. 15 individual microglial cells/mouse and n = 4-5 per group. Scale bar: A1-A4: 5 µm.

### Single-cell heterogeneity reveals distinct subpopulations of microglia in the Tg2576 mouse model of AD

We analyzed scRNA-seq data from 6,202 cells harvested from cortex and hippocampus of 15-month-old female WT and Tg2576 mice treated with PMX205 or vehicle (water) for 12 weeks (**Fig. S3A**). We clustered cells using Seurat (**Fig. S3B**) and identified the major populations, including astrocytes, microglia, ependymal cells, endothelial cells, perivascular macrophages, pericytes, OPCs, and oligodendrocytes, based on expression of canonical markers (**Fig. 5A**). Despite enriching for astrocytes and microglia during scRNA-seq extraction (see Methods), we did not recover the full spectrum of astrocyte subpopulations. We focused subsequent analysis on microglial cells, for which we obtained representative subpopulations. We recovered 10 distinct populations of microglia characterized by expression of canonical marker Csf1r (**Fig. 5B**). We also recovered a subpopulation of perivascular macrophages (PVM), which expressed canonical markers Cd163, Pf4, Cd209f, Cbr2, and Mrc1 (**Fig S3C**). Microglial subpopulations express distinct levels of homeostatic gene Tmem119 or disease-associated genes Cst7 and Lpl (**Fig. 5B**). These results emphasize the complexity and wide range of microglial response to amyloid pathology and PMX205 treatment.

**Fig 5.**
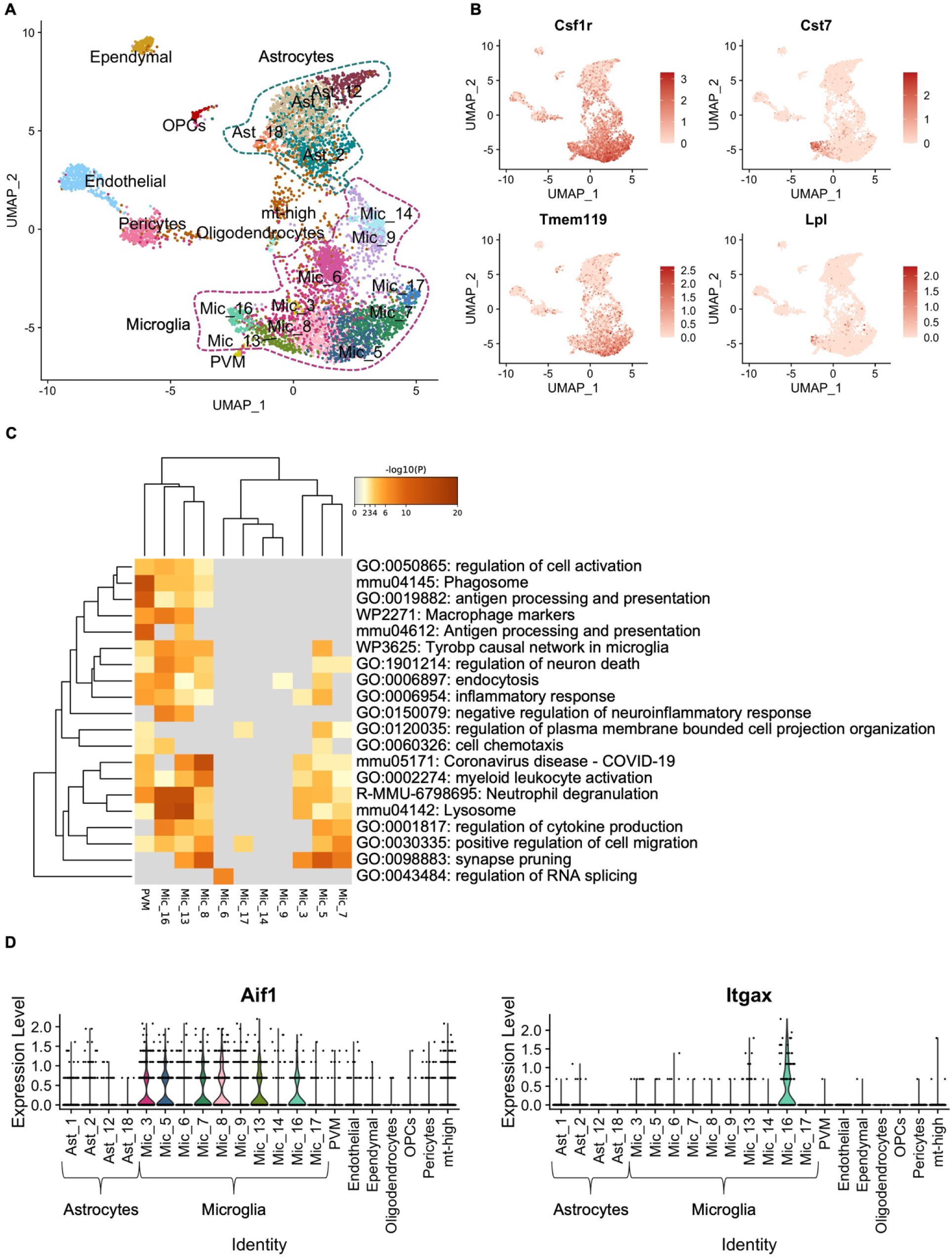
Single-cell RNA-seq reveals the complexity of microglial response to amyloid pathology. **(A)** UMAP embedding representation of 6,202 single-cells harvested from cortices and hippocampi of 15-month-old female Tg2576 and wild type mice. UMAP is annotated by clusters of major subpopulations recovered. **(B)** Distribution of the expression of signature homeostatic microglial genes Csf1r and Tmem119 and disease-associated microglial genes Cst7 and Lpl. **(C)** Gene ontology (GO) analyses of top 50 genes enriched in microglial and perivascular macrophage clusters. **(D)** Violin plot displays the expression levels of Aif1 (Iba1, left panel) and Itgax (Cd11c, right panel) per subpopulation.

We used Metascape’s multiple gene list feature to preform Gene ontology (GO) analysis of the top 50 markers per microglial cluster. GO analysis revealed that subpopulations 8, 13 and 16 are involved in cell activation, phagosome, and lysosome pathways (**Fig. 5C**), all of which are enriched in disease associated microglia (DAM) (*37, 38*). Expression of Aif1 (Iba1) was consistent across most microglial clusters (**Fig. 5D**, left panel), with Cx3Cr1, Csfr1 and Tmem119 expression also prominent within these clusters. Expression of Itgax was increased in microglia 16 (**Fig. 5D**, right panel). Interestingly, microglia 16 showed higher expression of Cst7 and Lpl (**Fig. 5B**).

### A microglial subpopulation enriched for synaptic pruning genes is reduced with PMX205 treatment

Examining the contribution of wild type versus Tg2576 cohorts to each group of cells, we observed that clusters 13 and 16 are mainly present in the Tg2576 mice (**Fig. 6A**), We further investigated the main pathways and biological processes enriched in the top 50 markers of clusters 8, 13, and 16, which showed relevant DAM signature GO terms, (**Fig. 6B**) and emphasizing that DAM microglia are enriched in the Tg2576 mice compared to WT, as expected in a mouse model of AD. Microglia 8, which express a distinct combination of homeostatic, DAM1 and DAM2 genes (**Fig. 6C**) is associated with synapse pruning. Notably, cluster 8 is reduced in the hippocampus of Tg2576 mice treated with PMX205, correlating with the reduced engulfment of synapses seen in our model (**Fig. 4 A5**). Our results suggest that PMX205 treatment reduces the activation of a unique microglial subpopulation associated with synapse pruning in the hippocampus of Tg2576 female mice and thus may reduce the neurodegenerative effects via reduction of excessive synapse pruning.

**Fig 6.**
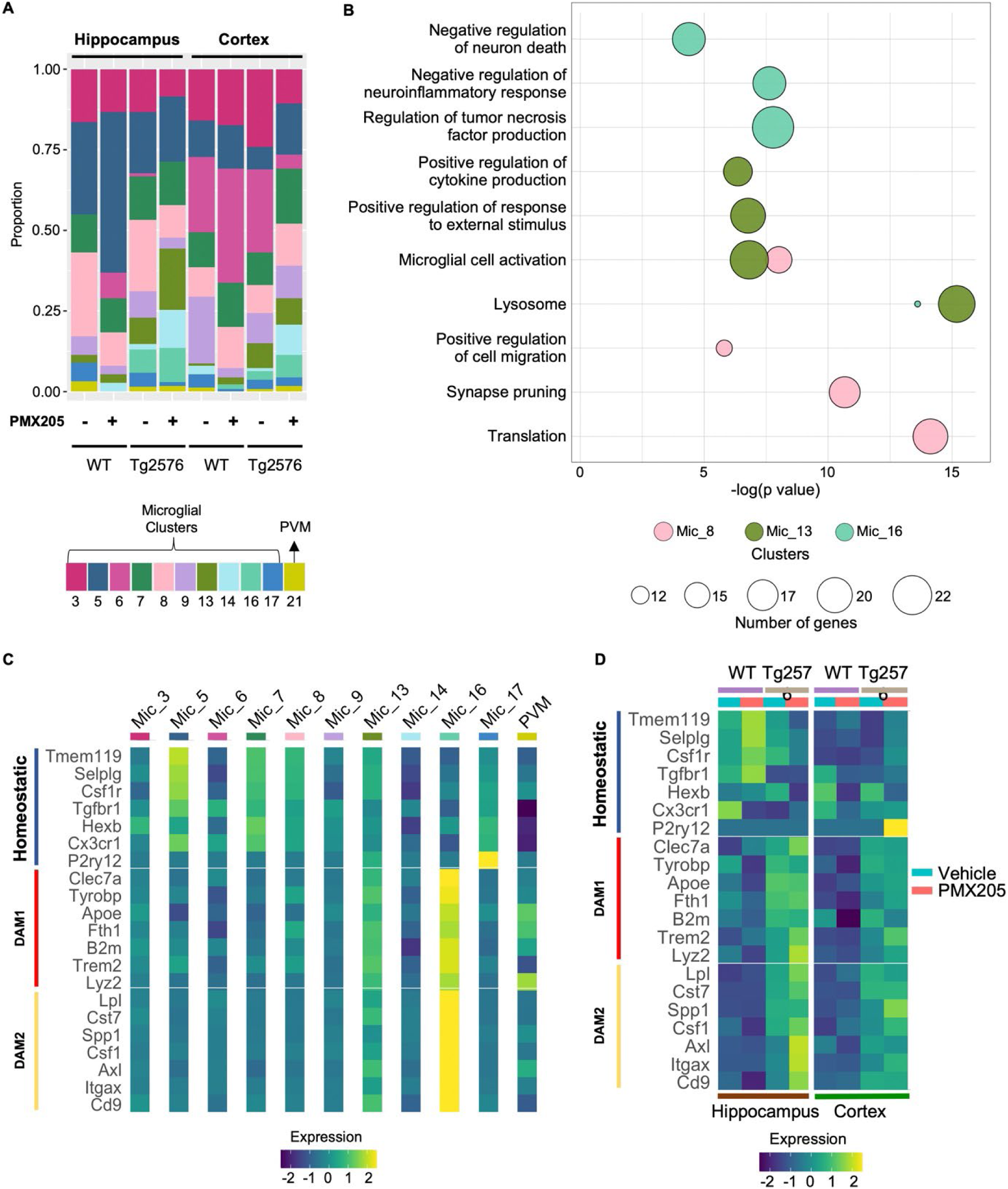
Changes in microglial identities post PMX205 treatment. **(A)** Bar plot of the relative proportion of microglial and perivascular macrophage subtypes per genotype and treatment. **(B)** GO Bubble plot of select top pathways and biological processes enriched in microglial clusters 8, 13 and 16. Circle size represents the number of genes enriched in each GO term. **(C-D)** Heatmap of signature homeostatic, DAM1 and DAM2 genes per microglial and PVM clusters. Normalized mean expression per cluster **(C).** Normalized mean expression per tissue, genotype, and treatment **(D).**

### PMX205 treatment up-regulates DAM2 microglial genes in the Tg2576 mouse model

Microglia 13, whose top markers include DAM1 signature genes, such as Lyz2, Trem2, Tyrobp, and Clec7a (**Fig. S3D**) are enriched in lysosome, microglial cell activation, and positive regulation of cytokine production GO terms. Microglia 16 overexpresses DAM2 genes, such as Lpl, Cst7, Csf1, and Itgax as well as genes such as GRN and Gpnmb that are involved in negative regulation of inflammatory response and negative regulation of neuron death. Both increased neuroinflammation and increased neuronal death can lead to worsening of AD pathology (*10, 39–41*) . Interestingly, the relative proportion of cluster 16 increases in the hippocampus (and cortex) of Tg2576 mice treated with PMX205, compared to vehicle (**Fig 6A**). Moreover, the levels of DAM2 genes increase with PMX205 treatment (**Fig. 6D**). PMX205 treatment also up-regulates homeostatic microglial genes in the WT mice. These findings suggest that PMX205 treatment reduces excessive inflammation and neuronal death in the Tg2576 mice and increases the activation of DAM2.

### App-CD74 signaling enriched in the Tg2576 mice treated with PMX205 may lead to reduced plaque load post-treatment

Intercellular communication is critical for response to inflammation, and apoptosis. To elucidate how PMX205 treatment affects microglia-microglia interactions in the Tg2576 mice we used the CellChat package. PMX205 treatment increased the global number of interactions between microglial subpopulations (**Fig. 7A**). Specifically, it increased the number of interactions within homeostatic microglia (Clusters 3, 5, 6, and 7) (**Fig. 7B**), as well as increased the signaling sent from homeostatic microglia, from cluster 16 (DAM2-like), and from PVM to all other microglial clusters. Next, we compared the signaling pathways identified by CellChat as significant in the Tg2576 mice treated with PMX205 or vehicle. Signaling pathways colored in coral are enriched in vehicle treated and pathways colored in turquoise are enriched in PMX205 treated mice. Incoming signaling from microglia 8 recovered from vehicle treated Tg2676 mice was enriched in “PROS” pathway (**Fig. 7C**), which corresponds to the ligation of Pros1 to Axl (**Fig. 7F**) in microglia. Vehicle treated microglia 13 was enriched for CCL pathway, which corresponds to the ligation of Ccl3 to Ccr5 (**Fig. 7D** **and 6F**). An excess of Ccl3 has been shown to impair spatial memory in mice (*42*). Ccl4 signaling sent from microglia 16 is enriched in PMX205 treated mice. Ccl4 stimulation has led to brain barrier disruption *in vivo* and *in vitro* (*43*), which can lead to increased PVM macrophage infiltration. Interestingly, in the PMX205 treated mice, both microglia 13 and 16 are sources of App signaling, which ligates to Cd74 receptors in PVM macrophages (**Fig. 7D-7F**). The interaction of CD74 and APP has been shown to suppress amyloid beta production *in vitro* (*44*). Our results suggest that PMX205 increase APP-CD74 signaling, which may help reduce the plaque load observed in the initial stages of the pathology in Tg2576 (**Sup Fig. 2 B1-B4 and C1-C4**).

**Fig 7.**
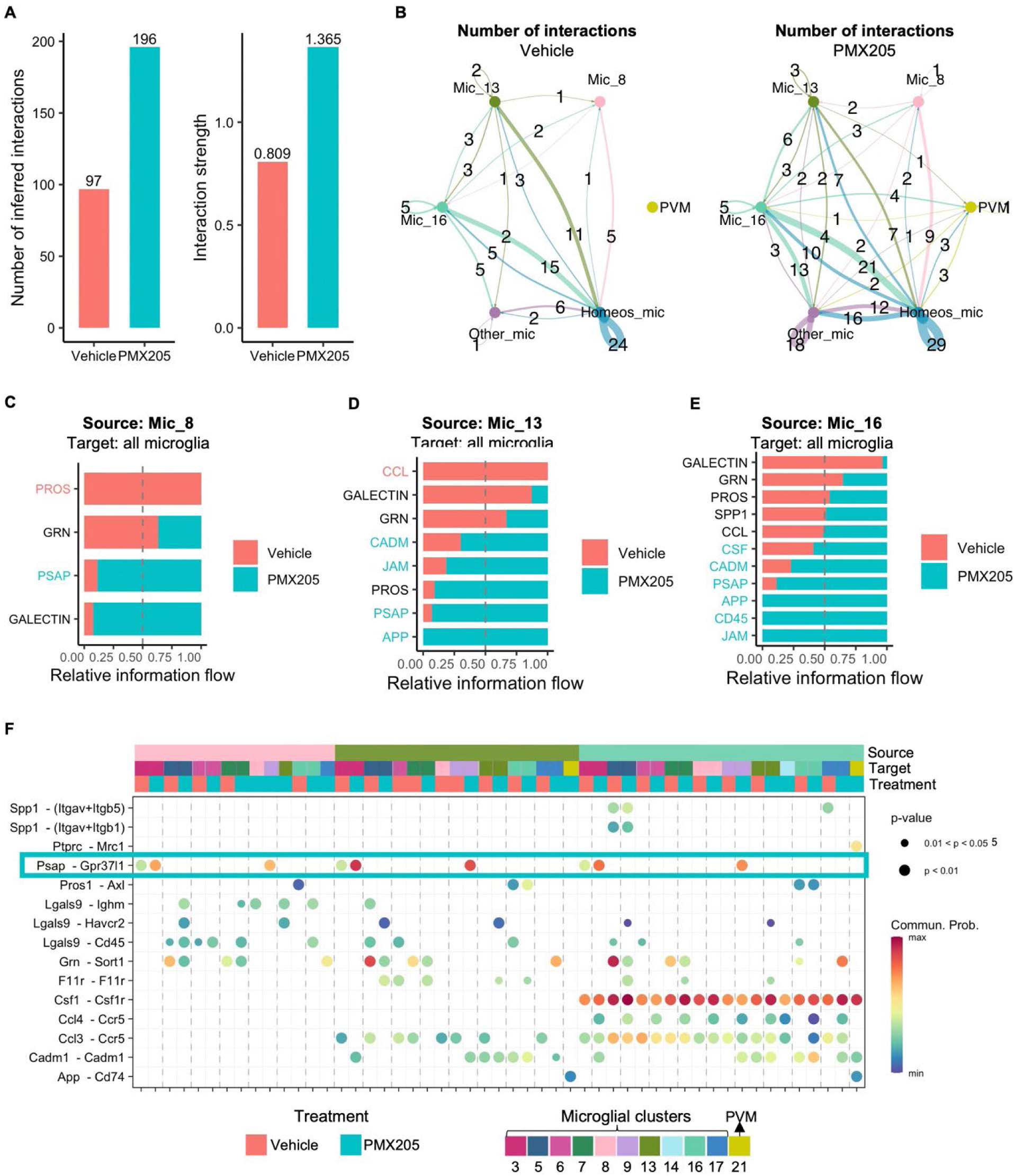
Increased microglia intercellular signaling in the Tg2576 mice treated with PMX205. **(A)** Comparison of the total number of microglia-microglia interactions and interaction strength per condition. **(B)** Aggregated cell-cell communication network among signature homeostatic microglial clusters, PVM, disease associated microglial clusters 8, 13 and 16 and other microglia. Thickness of the line is proportional to the number of interactions, which is also displayed. Circle size is proportional to the number of cells in each subpopulation. **(C-E)** Comparison of conserved and context specific signaling pathways ranked based on significance per condition. The top signaling pathways colored coral are enriched in vehicle treated Tg2576 mice and the ones colored turquoise are enriched in the PMX205 treated. Microglial clusters 8 **(C),** 13 **(D)** and 16 **(E)** are the sources of signaling to other microglia and PVM. **(F)** Communication probabilities mediated by ligand-receptors pairs. Signaling incoming from microglial clusters 8, 13 and 16 to other microglial and PVM subpopulations. Horizontal color bars indicate sources of signaling (top bar), targets (middle bar) and treatment (bottom bar). Vertical color bar indicates probability of interaction. Circle size indicates significance of interaction.

### Neuroprotective receptor GPR37L1 signaling is upregulated with PMX205 treatment

PMX205 treated microglia 16 send Csf1 signaling to all microglia subpopulations, consistent with a DAM2-like signature (**Fig. 7F**). This subpopulation is also enriched in CD45 signaling pathway (**Fig. 7E**). CD45 has been shown to be relevant for microglial clearance of oligomeric amyloid beta plaques in AD mouse models (*45*). Notably, in the presence of PMX205, microglia 8, 13 and 16 are enriched in the PSAP signaling pathway, which corresponds to the ligation of Psap (prosaposin) to Gpr37l1 (**Fig 7F**, turquoise rectangle). Prosaposin activation of receptor Gpr37l1 was found to be neuroprotective and glioprotective (*46*). This suggests that PMX205 treatment stimulates prosaposin signaling outgoing from activated microglia, which exerts neuroprotective functions in the Tg2576 mice.

## DISCUSSION

A role for the complement system during Alzheimer’s disease pathogenesis has been extensively demonstrated during the past 20 years by us and many others (reviewed in(*26*)). However, the specific consequences of C5a-C5aR1 modulation in the Alzheimer’s brain is not yet fully understood. The anaphylatoxin C5a is a potent inflammatory effector secreted after the cleavage of C5 as a downstream consequence of complement activation, that promotes chemotaxis and a strong inflammatory response when it binds to its receptor C5aR1 (reviewed in(*9*)). Up-regulation of the C5a receptor 1 during Alzheimer’s disease pathogenesis has been previously described in different mouse models (*19, 47*). Our previous studies have demonstrated a beneficial effect of the genetic ablation of C5aR1 in preventing loss of neurite complexity and cognitive behavior in the Arctic mouse model of AD (*18*). Further support of a protective role of a C5aR1 antagonist is the rescue of the excessive synaptic loss and decreased AD-like pathology in 2 other mouse models of AD(*20*) that correlates with memory loss and cognitive decline parameters seen in human AD(*32*). We recently described that C5a overexpression accelerated memory deficits while delaying the increase of several inflammatory genes until a later stage of the pathology in the Arctic mice (*19*). The latter effect is likely due to the abnormal elevated amounts of C5a (produced under the GFAP promoter) binding to C5aR2, which has been shown to possess anti-inflammatory properties(*48*), thus pointing to the benefit of specific inhibition of C5aR1 as a potential therapeutic target for Alzheimer’s disease.

Aß plaques can occur up to 20 years before the detection of cognitive decline in AD patients(*49, 50*). However, the mechanism by which amyloid plaques are formed, and more importantly, what is the role of microglial cells in plaque formation/elimination is still under debate. Studies point to a dual role of microglia in plaque formation and growth. Microglial depletion in the brain of the 5xFAD mouse model of AD at early ages resulted in almost no plaque formation(*51*), but removal of microglial cells in the aged brain (after plaques have accumulated) protects against synapse and neuronal loss with no effect on Aß plaques(*52*). On the other hand, in vivo studies by two-photon imaging in the same AD mouse model revealed that microglial death following Aß phagocytosis was a major contributor to plaque growth(*53*). Moreover, another study showed microglial cells around amyloid plaques phagocytose Aß via TAM receptors and convert the amyloid into a much more compacted material, adding to the growth of dense-core amyloid plaques(*54*). Reduced levels of microglial cells together with an impairment in Aß phagocytosis resulted in lower plaques numbers(*54*). These observations corroborate our earlier studies, which showed a significant reduction in several microglial activation markers accompanied by a profound decrease in amyloid plaques when the Tg2576 mouse model of AD was treated with a C5aR1 antagonist. However, here our Aß phagocytosis analysis did not reveal any significant changes when PMX205 was administer to the mice. We observed a reduction in the total number of amyloid plaques as well as reduced total Aß load, although the average size of the plaques present remained unchanged independently of the presence of C5aR1 antagonist. This suggests that C5a-C5aR1 signaling might influence the initial stage of seeding and formation of the Aß plaques but once these deposits are formed, their size keeps increasing with time and disease progression. Further studies are needed to elucidate the exact mechanism by which C5a-C5aR1 signaling influences amyloid plaque deposition during Alzheimer’s disease pathogenesis. It could be hypothesized that PMX205 might be directly interacting with Aß and altering fibrils formation; however, our previous in vitro studies reported no evidence of PMX205 affecting Aß fibrilization processes(*20*). Additionally, we previously reported a protective effect on neuronal complexity as well as in cognitive function when C5aR1 is genetically ablated in the Arctic48 mouse model(*18*). In this model, ablation of C5aR1 did not affect amyloid deposition; likely due to the Arctic48 mutation which is known to accelerate the Aß fibrilization process and results in fibrils highly resistant to clearance. This might explain the differences observed in plaque accumulation between the two mouse models(*18*).

Given the striking effect of C5aR1 inhibition on amyloid plaques in the Tg2576 mice, we also assessed the appearance of plaque associated dystrophic neurites. Dystrophic neurites are abnormal neuronal processes that appear associated with amyloid plaques in Alzheimer’s disease brains and can be considered as a sign of neurodegeneration. Recently, several authors hypothesized that microglial cells could form a physical barrier around amyloid plaques to protect from dystrophic neurite formation due to the amyloid toxicity(*55, 56*). Here, we reported a decreased in the total number of dystrophic neurites after PMX205 treatment; however, the ratio of dystrophic neurites per plaque did not change in response to C5aR1 inhibition, suggesting that once formed, amyloid plaques exert the same neurotoxicity to the surrounding neurites with or without C5aR1 inhibition. Since we also reported no significant changes in the halo of microglial cells around amyloid plaques in response to C5aR1 antagonist, we could presume that the beneficial effect in preventing the appearance of dystrophic neurites in response to PMX205 is mainly due to its impact (decrease) in the amount of amyloid pathology in the Tg2576 mouse model of AD.

Synaptic loss is one of the hallmarks of Alzheimer’s disease and its correlation with cognitive impairment and memory loss has been well described (*32, 57*). The role of complement-mediated microglial synaptic pruning during neurodegenerative diseases has been the focus of numerous studies during the past decade (reviewed in (*9*)). Among all the advances in the knowledge of the mechanism driving complement mediated synaptic pruning, we should highlight recent work showing that inhibition of C1q or C3 can successfully reduce the excessive synaptic loss that occurs with AD progression(*14, 33*). However, a complete inhibition of the complement cascade might not be a good therapeutic target for AD given the multiple beneficial roles of the upstream components of the complement system. In addition, such inhibition of C1q or C3 would also completely prevent downstream complement activation mediated events. In this manuscript, the treatment with a C5aR1 antagonist in the Tg2576 mouse model of AD significantly reduced the excessive microglial synaptic pruning in the CA3 hippocampal region, which in turn partially rescued the excessive pre-synaptic loss observed in vehicle treated AD mice (compared to WT littermates). These results reinforce findings from our previous work, in which we demonstrated that PMX205 resulted in a protective effect in hippocampal synaptophysin pre-synapses (*20*). In addition, by using high resolution confocal microscopy, we were able to image and quantify the number of VGlut1 synapses that were being actively engulfed by individual microglial cells. Furthermore, our scRNA-seq results identified a subset of microglial cells (cluster 8) that is directly involved in synaptic pruning processes. Interestingly, cluster 8 is downregulated in the hippocampus of Tg2576 animals treated with PMX205. Among the genes involved in this synaptic pruning GO biological process we found C1qa, C1qb, C1qc, Trem2 or C5aR1. Together, these findings suggest not only that C5aR1 could play an important role in the regulation of complement mediated microglial synaptic pruning but also that synaptic engulfment is a specialized function carried out by a small subset of microglial cells. Thus, the decrease in synaptic pruning that we observed is likely one of the mechanisms by which inhibition of C5a-C5aR1 signaling protects against cognitive deficits in AD mice.

The important role of microglia in AD pathology is well recognized. However, their molecular heterogeneity and diversity of functions during disease progression are only now being defined. Recent studies support both beneficial, as well as detrimental roles for microglia in Alzheimer’s disease (*58*). Disease associated microglia have been reported to form two distinct subpopulations, termed DAM1 which overexpress B2m, Tyrobp, and Apoe, as well as DAM2, which overexpress Spp1, Itgax and Axl (*59, 60*). Our Gene Ontology analysis revealed that Microglial 16 (considered as a DAM2 subpopulation) is involved in the negative regulation of inflammatory response, involving genes such as Cst7, Ccl3, Ccl4, Grn or Gpnmb, among others. While the specific role of those genes in regulating the inflammatory response is still not clear, Kang et al suggested that Ccl3 and Ccl4 might be crucial for microglial function in aging and disease, potentially acting through the recruitment of peripheral immune cells(*61*). While further analysis is necessary due to low numbers of cells in this cluster, a consistent hypothesis is that Ccl3 and Ccl4 might be attracting PVM macrophages that express CD74 receptors, while detrimental responses due to C5a-C5aR1 would be prevented due to PMX205. As we mentioned above, our CellChat results showed an enhancement of APP-CD74 signaling when PMX205 was administer to Tg2576 mice, which could exert a beneficial effect in reducing the amyloid pathology. Moreover, in a mouse model of AD, the loss of Grn has been shown to increase plaque load and exacerbate cognitive deficits, while its overexpression exerted a protective effect in neuronal loss and spatial memory(*62, 63*). Microglia which upregulate Trem2 seem to be crucial to help clear up Aβ plaques (*64*). Thus, Trem2 enables protective microglial responses during AD pathogenesis (*65*). Our results show that a subtype of microglia that exhibits a DAM-like signature is enriched in the hippocampus of Tg2576 mice following PMX205 treatment. We also observed overexpression of Trem2 post treatment, suggesting that the strong reduction in the number of cortical ThioS+ plaques we observed is linked to an increase in DAM population with disease mitigating effects.

Prosaposin (PSAP) is a multifunctional protein that regulates lysosomal enzymes intracellularly and also acts as a neuroprotective secreted factor extracellularly (*66*). Patients lacking prosaposin presented increase in reactive astrocytes and lack of mature oligodendrocytes and cortical neurons (*67, 68*). The Psap-Gpr37l1 signaling has been described to induce production of neurotropic factors by microglia for neuronal recovery (*69*). Our single cell signaling inference results suggest that PMX205 treatment increases prosaposin signaling secreted from disease associated microglia targeting other microglial subtypes that express Gpr37l1. Ligation of Psap to Gpr37l1 might be one of the neuroprotective pathways stimulated by PMX205 treatment.

Our study has some limitations. First, due to the technology we used for single cell RNAseq, we isolated microglial cells and did not analyze possible changes in the astrocytic population. Since it is known that astrocytes also play a key role during Alzheimer’s disease pathogenesis, our future studies will investigate the potential effect of PMX205 into astrocytic subpopulations. Another limitation is that our amyloid phagocytosis analysis was done in fixed tissue and so, we could not evaluate potential lysosomal dysfunction or abnormalities in AD mice. 2-photon in vivo studies can be used to overcome this limitation in future studies. Nonetheless, our findings have important implications regarding the role of microglial cells during AD pathogenesis and point out to the antagonism of C5aR1 as a possible AD therapeutic strategy.

In summary, here, we provide evidence of a neuroprotective effect of the pharmacological inhibition of C5a-C5aR1 signaling in the pathology progression in the Tg2576 mouse model of Alzheimer’s disease. PMX205, a C5aR1 antagonist, leads to significant reduction in the amyloid burden as well as slows the appearance of plaque-associated dystrophic neurites. Furthermore, this is the first study that shows the single cell heterogeneity of an AD mouse model following treatment with PMX205. We identify specific microglial subpopulation changes and signaling pathways that are consistent with being key mediators of the beneficial effects observed after this treatment. We report a microglial subtype involved in synapse pruning, which is reduced with PMX205 treatment, as well as an increase in hippocampal DAM2 microglia. Moreover, we identify a possible increase in prosaposin signaling and Gpr37l1 activation following PMX205 treatment. Importantly, our results point to PMX205 as a promising narrowly targeted therapeutic strategy to treat Alzheimer’s disease.

## MATERIALS AND METHODS

### Study design

The studies described in this manuscript were conducted to further characterize the effect of a C5aR1 antagonist (PMX205) on microglial subpopulations in a mouse model of Alzheimer’s disease (Tg2576). To address this question, we used Tg2576 and WT littermates and treated them either with PMX205 or regular drinking water as a control for 12 weeks. Using immunohistochemistry and single cell RNAseq analysis, we evaluate the effect of blocking C5a-C5aR1 signaling at an early stage of Alzheimer’s disease (at the onset of amyloid pathology). The sample size for animal experiments used here was based on our previous studies and we used the minimum number of animals needed to obtain significant results. All animal studies were approved by The Institutional Animal Care and Use Committee of University of California at Irvine.

### Animals

All animal procedures were approved by The Institutional Animal Care and Use Committee of University of California at Irvine and experiments were performed according to the NIH Guide for the Care and Use of laboratory animals. Mice were single housed during drug treatment starting at 12 mo of age and given access to food and water ad libitum. Tg2576 mice, developed by K. Hsiao(*70*), overexpressed the 695 isoform of the amyloid precursor protein (APP) with the Swedish mutation (KM670/671NL) under the control of the prion promoter on a B6/SJL genetic background. Only female hemizygous Tg2576 mice were used as they develop cortical amyloid plaques by 11-13 months of age, earlier than male Tg2576(*70*). WT (B6/SJL) female littermates were used as control mice.

### C5aR1 antagonist (PMX205) treatment

C5aR1 antagonist, PMX205, was kindly provided by Dr. Ian Campbell, Teva Pharmaceuticals, West Chester, PA. Treatment with PMX205 was administered in the drinking water at 20 μg/ml to mice; the starting point was before the onset of amyloid pathology at 12 months old for the Tg2576 mice (and B6/SJL WT littermates) (see Sup Figure 1A). 12-months old mice were divided into 4 groups: WT-vehicle, WT-PMX205, Tg2576-vehicle and Tg2576-PMX205. PMX205 treatment was administer for 12 weeks (equivalent to ~ 2-8 mg/kg/day), based on previous results from our lab(*20*). Mice were singly housed and had free access to the drinking water during the whole treatment. Mouse weight and volume of PMX205/vehicle consumed by each mouse were measured weekly.

### Tissue preparation

At the end of the PMX205/vehicle treatment mice (n=4-5/group) were deeply anesthetized with isoflurane and transcardially perfused with HBSS modified buffer (Hank’s Balanced Salt Solution without Calcium and Magnesium), containing Actinomycin D (5 μg/ml) and Triptolide (10 μM)(*71*). For immunohistochemistry, half brains were fixed in 4% paraformaldehyde for 24 hours, sectioned at 40 μm thickness in the coronal plane on a vibratome (Leica VT1000S) and then stored in PBS-0.02% sodium azide at 4°C. For scRNAseq, half brains were processed as detailed below.

### Antibodies

The following primary antibodies were used for this study: rabbit polyclonal anti-oligomeric Aβ OC (1:1000, Millipore #AB2286), mouse monoclonal anti-Aβ 6E10 (1:1000, Biolegend #A03001), rabbit polyclonal anti-Iba1 (1:1000, Wako #019-19741), rat monoclonal anti-CD68 (1:700 Biolegend #137001), rat monoclonal anti-CD11b (Biorad #MCA74G), Armenian hamster monoclonal anti-CD11c (1:400, Biorad #MCA1369), rat monoclonal anti-C3 (1:50, Hycult #HM1045), rabbit polyclonal anti-GFAP (1:3000, Dako #Z0334), chicken polyclonal anti-GFAP (1:1000, Abcam #ab4674), rat monoclonal anti-LAMP1 (1:500, Abcam #ab25245), guinea pig polyclonal anti-VGlut1 (1:1000, Millipore #AB5905). Alexa Fluor secondary antibodies were diluted to 1:500 in blocking solution and included: 488 goat anti-rat (Invitrogen #A21212), 488 goat anti-chicken (Invitrogen #A11039), 488 goat anti-guinea pig (Invitrogen #A11073), 488 goat anti-mouse (Invitrogen #A11001), 555 goat anti-rabbit (Invitrogen #A21428), 555 goat anti-rat (Invitrogen #A21434), 568 Goat anti-Armenian hamster (Abcam, # ab175716), 647 goat anti-rabbit (Invitrogen #A21244).

### Immunofluorescence (IF)

Free-floating sections were briefly washed with 0.1M PBS. For general antigen retrieval method, brain sections were incubated in 50 mM citrate buffer pH 6.0 for 30 min at 80 °C. Sections were then incubated in blocking solution (0.1M PBS, 2% BSA, 10% NGS and 0.1% Triton X100) for 1 hour at room temperature (RT) followed by incubation with primary antibodies in blocking buffer for 24h at RT. After washing, corresponding secondary antibodies in blocking buffer were added for 1h at RT. To counterstain with Thioflavin-S (Sigma-Aldrich #T1892), sections were incubated in 0.1% ThioS diluted in MilliQH_2_O for 10 minutes after the secondary antibody and washed with 50% ethanol followed by PBS. Amylo-Glo (1:100, Biosensis #TR-300-AG) staining was performed before the initial blocking step. Briefly, tissues were incubated in Amylo-Glo for 10 min at RT, then washed with PBS and rinsed with ddH_2_O. Sections were then mounted onto a slide and coverslipped with Hardset Antifade Mounting Medium (Vectashield, Vector). 3-4 sections/mouse were stained and processed at the same time, using batch solutions, and imaged via a Slidescanner (Zeiss Axio Scan.Z1) using a 10x objective. In addition, higher magnification images were taken under a confocal microscope (Leica SP8) using a 63x objective. For 3D reconstructions, serial confocal images (z-stacks) of Iba1/CD68/Vglut1 and Iba1/CD68/6E10 were acquired in 0.3 μm or 0.5 μm per step, respectively, under a 63x Leica SP8 confocal microscope.

### Imaris Quantitative Analysis

Image analyses were carried out in the whole hippocampus and cortex, using Imaris 9.7 software (Bitplane Inc.). ThioS, OC, Iba1, CD68, Cd11b, Cd11c, GFAP, C3, and Lamp1 immunopositive signals within the selected brain region were identified by a threshold level mask, which was maintained throughout the whole image analysis for uniformity. Quantitative comparisons between groups were carried out on comparable brain sections processed at the same time with the same batches of solutions for uniformity. Amyloid burden was acquired by measuring both, the number and size (in area units, μm^2^) of amyloid plaques in the selected brain regions. Astroglial, microglial and dystrophic neurite load (Field Area %) were obtained by using the surface option from Imaris and normalized to the total area of the hippocampus or cortex. The number of microglia and astroglial cells was obtained with the spots option in Imaris and normalized to the total area of hippocampus or cortex. Synaptic engulfment was quantified in 15 microglial cells/mouse with 4-5 mice/group, and it was defined as the co-localization (defined at ≤ 200nm distance) of Vglut1+ synapses (detected by Imaris spots) with CD68 and Iba1 surfaces and normalized to the total volume of the image. Microglial phagocytosis of Aβ was analyzed in 15 amyloid plaques (randomly selected) per mouse per group and phagocytosis index was defined as the co-localization of Iba1, CD68 and 6E10 surfaces, normalized to the total amount of Aβ per image. Microglial engagement with amyloid plaques was also analyzed in 15 amyloid plaques (randomly selected) per mouse per group and quantified as the ratio of co-localization of Iba1 and 6E10 surfaces to the total Aβ.

### scRNA extraction

Half brains were quickly dissected into cortex and hippocampus on ice and dissected fresh tissue was then transferred to an inhibitor cocktail (HBSS with Actinomycin D (5 μg/ml), Triptolide (10 μM) and Anisomycin (27.1 μg/ml)) at 4°C to prevent microglial activation (*71*). Tissues were manually dissociated with a razor blade on ice and placed in the gentleMACS^TM^ Octo Dissociator and Adult Brain Dissociation Kit (Cat# 130-107-677) (Miltenyi Biotec), supplemented with the inhibitor cocktail, for enzymatic dissociation. Following dissociation, cells were filtered through 70 µl SmartStrainers (Cat# 130-110-916), debris was removed with debris removal solution provided in the Adult Brain Dissociation Kit, and myelin was cleared with Myelin removal beads II (Cat# 130-096-733). Microglia and astrocytes were respectively isolated using Anti-CD11b and Anti-ACSA-2 MicroBeads (Miltenyi Biotec). Cells were counted and resuspended in 10-15 µL and 20-30 µL of buffer (0.05% BSA in DPBS) for half hippocampus and half cortex, respectively, to reach a minimum of 1,000 cells/µL up to around 7,000 cells/µL.

### scRNA-seq library preparation and sequencing

Single cell libraries were built using the Illumina Bio-Rad SureCell WTA 3’ Library Prep Kit (Cat# 20014280). Isolated single cells were loaded into the Bio-Rad microfluidic device. Cells were lysed, barcoded, and reverse transcribed inside nanodroplets. The first strand cDNA was pooled from all the nanodroplets and the second strands were synthetized. The cDNA was cleaned, tagmented, and amplified to generate final libraries. Library concentration and size distribution were assessed using the Agilent 2100 Bioanalyzer. Library average length was 900-1100 base-pairs. Single-cell libraries were sequenced using a 150-cycle NextSeq 500/550 High Output Kit v2.5 (Cat# 20024907) on an Illumina NextSeq500 between 2.0-2.9pM loading concentration using a custom primer, and with each library having a minimum depth of 50 million reads.

### scRNA-seq processing and data analysis

Fastq files were processed using the KB_python pipeline, which is based on kallisto bustools using the parameters: kb count -x SURECELL –h5ad -t 20. Each cell was mapped to the mouse genome mm10 with annotations from GENCODE v21(*72, 73*). Low quality cells with less than 500UMI, more than 5% mitochondrial reads, and less than 300 genes detected were removed from downstream analysis. Doublets were removed using Scrublet(*74*). Normalization and clustering were calculated using Seurat SCT and Leiden algorithms, respectively(*75*). Intercellular communication between clusters of cells was inferred using CellChat(*76*).

### Statistical analyses

All immunofluorescent data were analyzed using either a Student’s t-test or two-way ANOVA, followed by Tukey’s post hoc test for comparisons among more than 2 groups, using GraphPad Prism Version 9 (La Jolla, CA). The significance was set at 95% of confidence. Linear correlations were analyzed using the Pearson’s test. *p<0.05, **p<0.01, ***p<0.001 and ****p<0.0001. Data were presented as the mean ± SEM (standard error of the mean).

## ACKNOWLEDGEMENTS

This study was made possible in part through access to the Optical Biology Core Facility of the Developmental Biology Center, a shared resource supported by the Cancer Center Support Grant (CA-62203) and Center for Complex Biological Systems Support Grant (GM-076516) at the University of California, Irvine.

## Author Contributions

Conceptualization: AGA, KC, GBD, AM, AJT

Funding acquisition: AM, AJT

Investigation: AG, KC, GBD, PS, NDS, SHC, HYL, MAP, TJP

Visualization: AG, KC

Writing – original draft: AG, KC

Writing – review and editing: AG, KC, GBD, PS, NDS, SHC, HYL, TJP, MAP, AM, AJT

## Data and materials availability

The accession number for the sequencing data reported in this paper is GEO: GSE200942.

## SUPPLEMENTARY MATERIALS

**Supplemental Figure 1:**
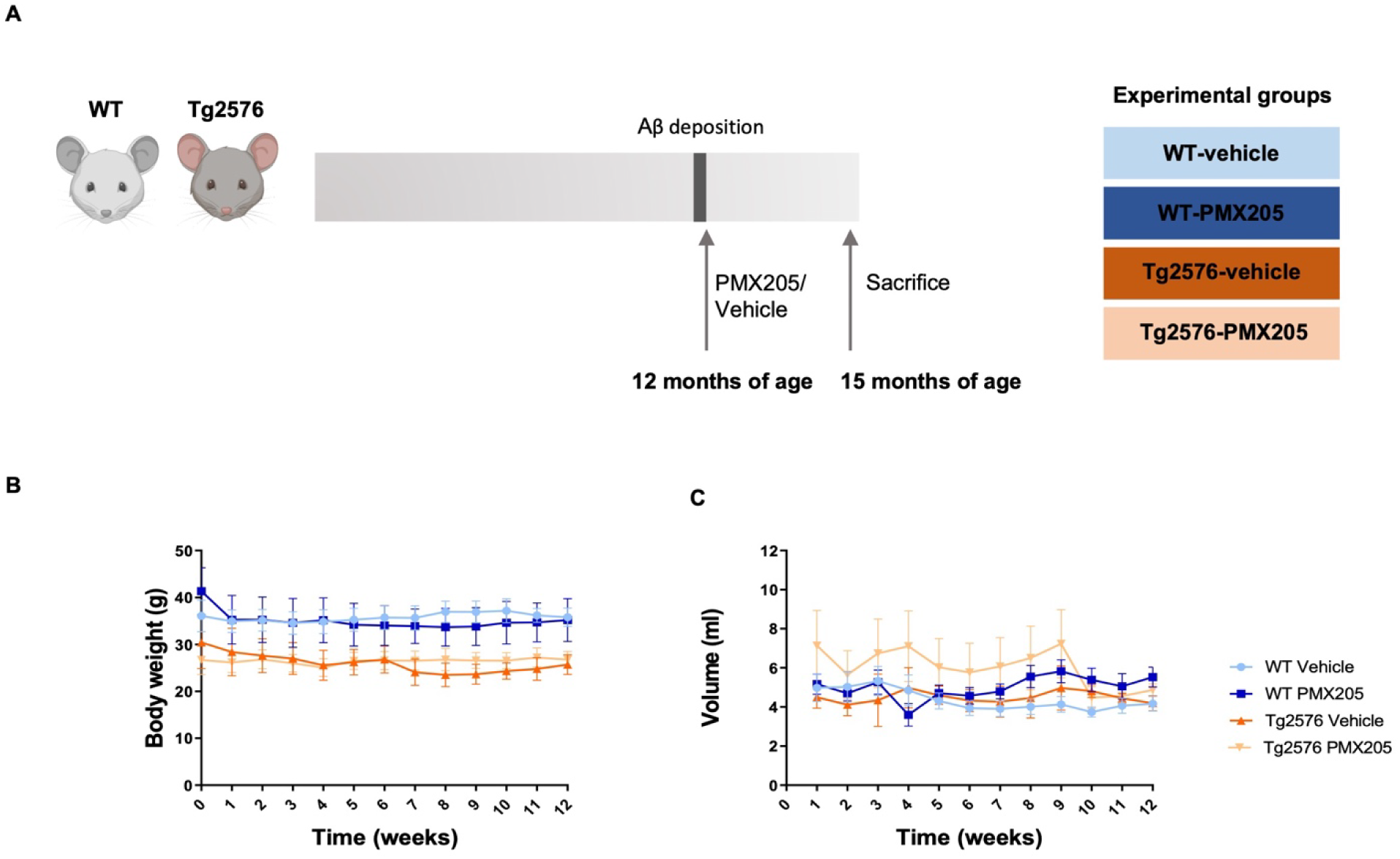
PMX205 treatment of Tg2576 and WT mice. (**A**) Schematic diagram of experimental design showing that Tg2576 mice and WT littermates were treated or not with 20 µg/ml of PMX205 for 12 weeks from 12-15 months of age. (**B-C**). During the duration of the treatment, mice body weight was recorded every week and no significant changes were observed in any of the experimental groups. Average drinking volume/week for each mouse showed no significant changes between the PMX205 and the control groups **(C).** Data shown as Mean ± SEM. Statistical analysis used a two-way ANOVA followed by Tukey’s post hoc test, n = 4-5 per group.

**Supplemental Figure 2:**
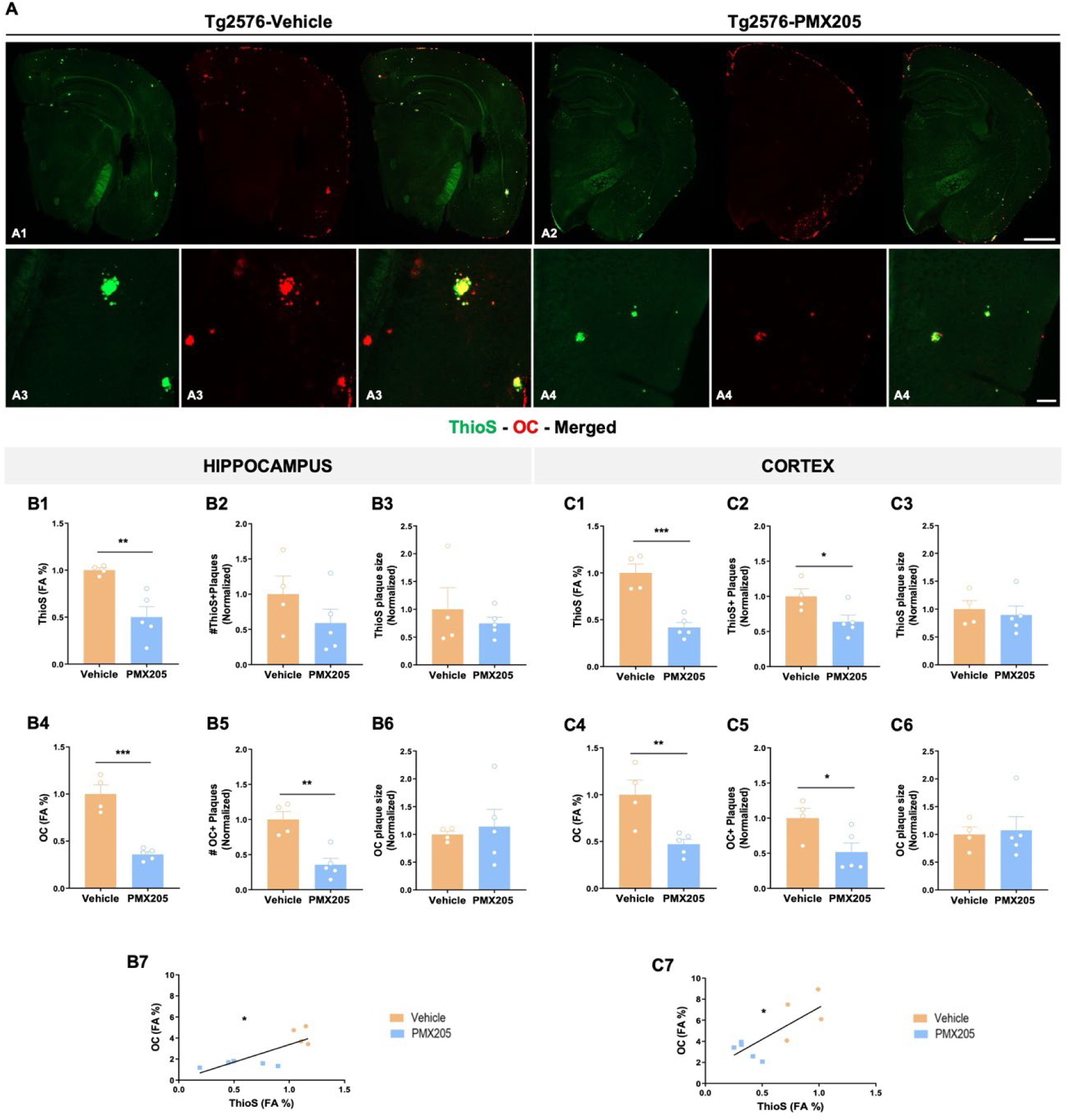
Amyloid pathology decrease with PMX205 in the Tg2576 mouse model of Alzheimer’s disease. (**A**) Representative stitched images of brain hemispheres of Tg2576-vehicle (**A1**) and Tg2576-PMX205 (**A2**) stained with ThioS, and OC. Inserts (**A3-A4**) shows higher magnification images of Aß deposition in the cortex. (**B-C**) Quantification of ThioS field area percent (FA%) showed a significant reduction of amyloid burden in the hippocampus (**B1**) and the cortex (**C1**). Similarly, OC FA% is strongly reduced in both regions analyzed (**B4** and **C4**). Quantification of ThioS+ and OC+ number of plaques (**B2**, **C2**, **B5** and **C5**) and size of plaques (**B3**, **C3**, **B6** and **C**6) revealed a reduction in the total number but not in the size of amyloid deposits. Positive correlation of ThioS FA% and OC FA% in both, hippocampus (R^2^ = 0.6), and cortex (R^2^ = 0.58) (**B**7 and **C7**). Data are shown as Mean ± SEM and normalized to control group (Tg2576-vehicle). Statistical analysis used a two-tailed t-test and Pearson’s correlation test. Significance is indicated as * p < 0.05; ** p < 0.01 and *** p < 0.001. 3 sections/mouse and n = 4-5 mice per group. Scale bar: A1-A2: 1000 µm; A3-A4: 100 µm.

**Supplemental Figure 3:**
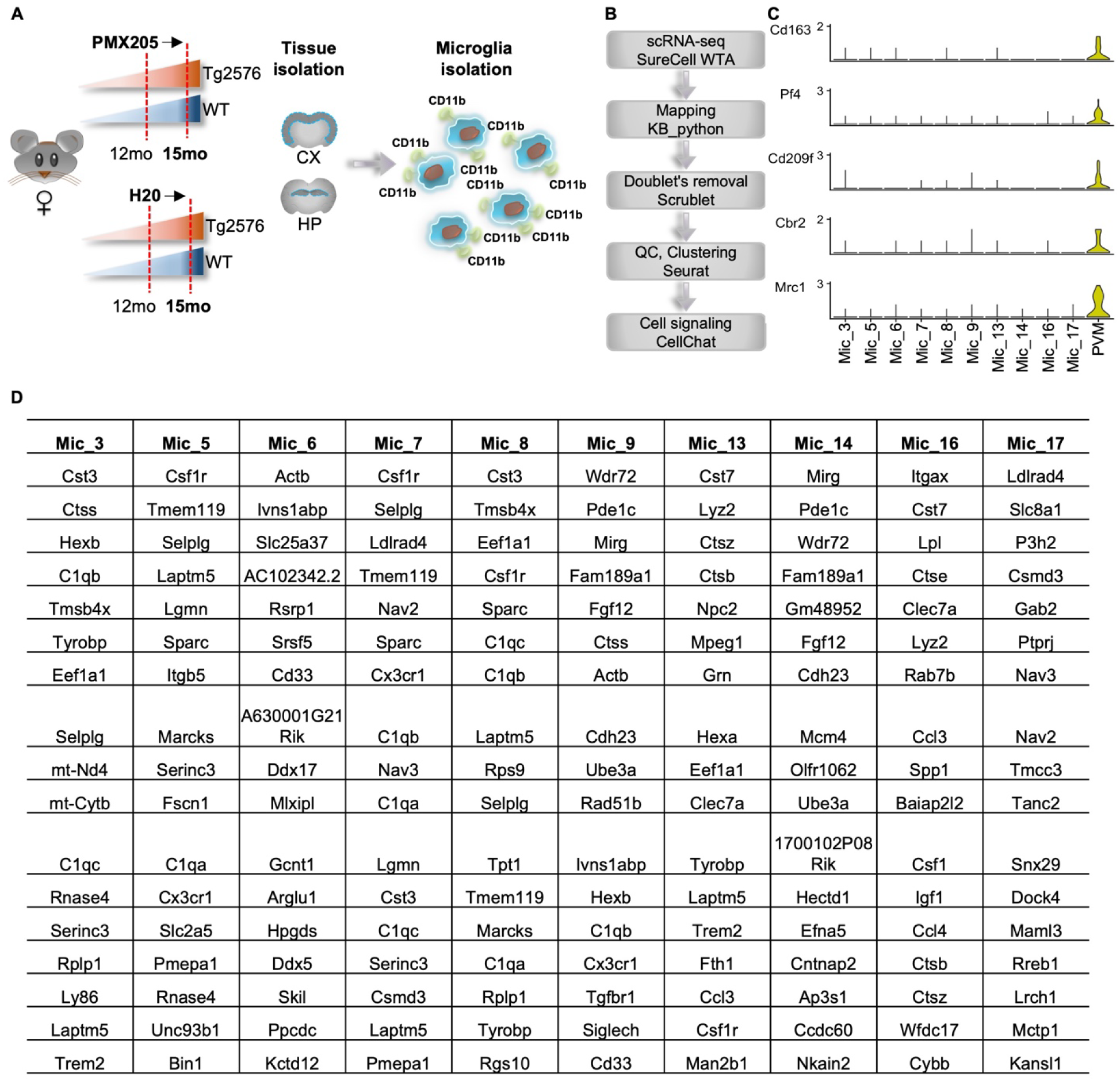
Single-cell RNA-seq of WT and Tg2576 female mice reveals distinct populations of macrophage and microglia. **(A)** Single-cell RNA-seq data collection pipeline. **(B)** Single-cell RNA-seq data analysis pipeline. **(C)** Expression of canonical perivascular macrophage markers across subpopulations. **(D)** Top 17 markers per microglial cluster (logFC >0.25, positive only).

